# Normal mammary gland development after MMTV-Cre mediated conditional PAK4 gene depletion

**DOI:** 10.1101/658583

**Authors:** Parisa Rabieifar, Ting Zhuang, Tânia D. F. Costa, Miao Zhao, Staffan Strömblad

## Abstract

p21-activated protein kinases (PAKs) are serine/threonine kinases functioning as downstream effectors of the small GTPases Rac1 and Cdc42. Members of the PAK family are overexpressed in human breast cancer, but their role in mammary gland development is not fully explored. Here we examined the functional role of PAK4 in mammary gland development by creating a mouse model of MMTV-Cre driven conditional PAK4 gene depletion in the mammary gland. The PAK4 conditional knock-out mice were born healthy with no observed developmental deficits. Mammary gland whole-mounts revealed no defects in ductal formation or elongation of the mammary tree through the fat pad. PAK4 gene depletion also did not alter proliferation and invasion of the mammary epithelium in young virgin mice. Moreover, adult mice gave birth to healthy pups with normal body weight upon weaning. This implies that MMTV-Cre induced gene depletion of PAK4 in mice does not impair normal mammary gland development and thereby provides an *in vivo* model for examination of the potential function of PAK4 in breast cancer.

## Introduction

The mammary gland is a highly ductal organ mainly composed of two distinct cell compartments, the epithelium, and the surrounding stroma, which are derived from ectoderm and mesoderm during embryogenesis ^1–3^. Unlike most other epithelial organs, development of the mammary gland occurs postnatally ^4–6^. Many signaling molecules such as growth hormone, estrogen, and growth factors stimulate formation and invasion of terminal end buds (TEB) to the mammary fat pad by regulating extracellular matrix (ECM) proteins ^5,7,8^.

Within the ECM, integrins act as chemomechanical sensors of mammary epithelial cells, which thereby receive signals from the surrounding environment and regulate cell proliferation, polarization, and further morphological organization through its downstream mediators such as the family of small Rho GTPases ^9,10^.

The p21-activated kinase (PAK) family of serine/threonine protein kinases are downstream effectors of the small Rho family GTPases Cdc42 and Rac ^11,12^. PAKs are categorized into two subgroups based on their sequence homology. Group I includes PAK1, 2 and 3, while the Group II includes PAK4, 5 and 6.

PAKs control major subcellular activities, such as cytoskeletal remodeling, mitotic progression, DNA damage response and play essential roles in organ formation throughout mammalian development ^13,14^. Moreover, their over-expression is associated with cell proliferation, cell survival, invasion, angiogenesis and epithelial-mesenchymal transition (EMT), which are all necessary for cancer initiation and progression ^12^.

PAK4, a member of group II PAKs is expressed both during development and in adult tissues ^15–18^. PAK4 is involved in a variety of cytoskeletal regulation such as promoting filopodia formation, dissolution of stress fibers, controlling actin polymerization and depolymerization as well as focal adhesion turnover^19–25^. Consistently, PAK4 overexpression is correlated with poor patient outcome in breast cancer patients, and its overexpression in breast cancer cell lines was shown to increase cell survival, anchorage-independent growth, cell migration and invasion^26–30^. PAK4 also plays an essential role during embryonic development, as complete depletion of PAK4 in mice caused embryonic lethality with severe defects in the heart, brain, and vasculature of the animals; however, its role in mammary gland development has not been investigated ^31–33^. Due to the early embryonic lethality of conventional PAK4 knock-out mice, conditional PAK4 knock-out mice have been developed to study its role in different tissue’s development ^16,33^. To this end, we created a transgenic mouse model to conditionally deplete PAK4 in the mammary gland by crossing MMTV-Cre mice with PAK4 ^floxed/floxed^ mice. We observed that PAK4 conditional knock-out mice were viable, produced standard litter sizes and pups with normal body weight upon weaning. Moreover, PAK4 knock-out mice exhibited healthy ductal morphology. These results indicate that PAK4 is dispensable for mouse mammary gland development.

## Results

### The spatial activity of Cre and disruption of PAK4 in the mouse mammary gland

Given that complete depletion of the PAK4 gene in the mouse causes embryonic lethality^16^, we created a mouse model to deplete PAK4 in the mammary epithelium using the Cre/loxP system; by crossing MMTV-Cre mice with PAK4-floxed mice ^31,33,34^. MMTV-Cre mice (line D) have been used in breeding, as this line has minimal effects on mammary gland development compared to other lines ^35^. The MMTV-Cre mice were used as a control group in this study (Fig.1 a). PAK4 flox was genotyped by PCR analysis of genomic DNA, identifying wild-type mice, as well as homozygous and heterozygous PAK4 knock-out mice (Fig.1 b). When MMTV-Cre; PAK4^fl/+^ mice were crossed with PAK4^fl/fl^ mice, four different genotypes were born approximately at the expected Mendelian ratio, i.e. Wild-type (PAK^fl/fl^ and PAK4^fl/+^) 49%; Het (MMTV-Cre; PAK4^fl/+^) 23%; and Homo PAK4 KO (MMTV-Cre; PAK4^fl/fl^) 28% (Table 1). Hereafter, for simplicity mice with MMTV-Cre; PAK4^fl/fl^ genotype will be referred as PAK4^MEp-/-^ and MMTV-Cre mice as PAK4^MEp+/+^. Within the same genotypes, female and male displayed an approximately equal distribution; suggesting that loss of PAK4 in the mammary epithelium does not affect survival in any of the sexes.

**Figure 1:**
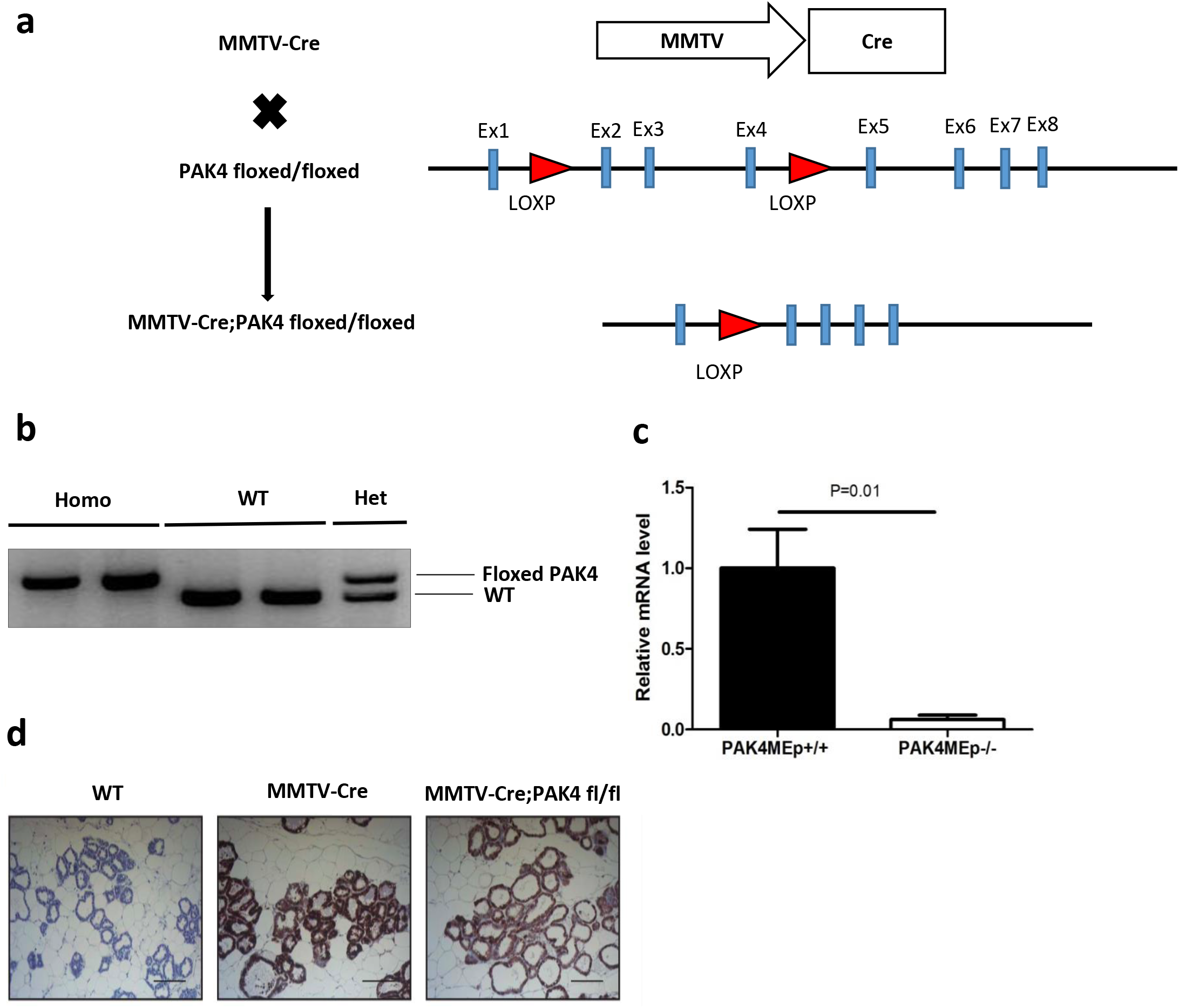
Conditional gene depletion of PAK4 in the mammary epithelium using the Cre-Lox system. **(a)** Graphical representation of the strategy for the generation of a mouse model with conditional PAK4 gene depletion in the mammary epithelium. Exons are indicated by light blue rectangles and Lox P sites are indicated by red triangles. As previously described Sox2-Cre expression and consequent recombination of LOXP sites result in the depletion of exons 2-4 in the mouse PAK4 gene as previously described ^33^. **(b)** PCR analysis of genomic DNA from 21 days old mouse. The upper band displays the floxed PAK4 allele, while the lower band displays a PAK4 WT allele. Thus, the appearance of the upper band alone displays homozygous floxed PAK4 allele; the lower band alone represents WT PAK4 allele; while the presence of both bands means that the mouse is PAK4 heterozygous, i.e. one WT and one floxed allele. Full gels are showed in supplementary Fig. S1a **(c)** PAK4 mRNA expression comparing PAK4^MEp-/-^ and PAK4^MEp+/+^ mice. mRNA was isolated from Lactation day 2 mice thoracic mammary glands and quantified by qRT-PCR. Shown values represent mean ± s.d. (n = 3 mice per group). P=0.01 for the PAK4^MEp-/-^ group versus PAK4^MEp+/+^ group, according to unpaired t-test. **(d)** Immunohistochemistry staining of Cre in thoracic mammary glands isolated from mouse at Lactation day 2 of different genotypes. Scale bar: 100 μm.

**Table 1:**
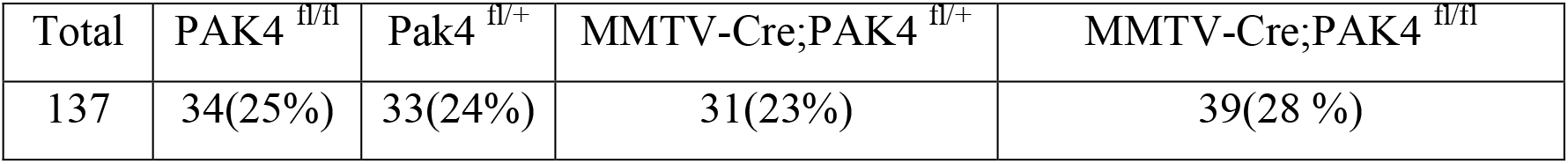
Genotypes of the progeny from MMTV-Cre; PAK4 fl/+ with PAK4 fl/fl

| Total | PAK4<sup>fl/fl</sup> | Pak4<sup>fl/+</sup> | MMTV-Cre;PAK4<sup>fl/+</sup> | MMTV-Cre;PAK4<sup>fl/fl</sup> |
| --- | --- | --- | --- | --- |
| 137 | 34(25%) | 33(24%) | 31(23%) | 39(28 %) |

To test the efficiency of the PAK4 gene depletion in PAK4^MEp-/-^ mouse mammary glands, mRNA levels of PAK4 was analyzed in thoracic mammary glands isolated from mice in the lactation day 2 stage. Importantly, we found a significant suppression of PAK4 mRNA expression in the mammary glands from PAK4^MEp-/-^ transgenic mice compared with mammary glands from matched PAK4^MEp+/+^ mice (Fig.1c). Because of the current lack of reliable PAK4 antibodies for immunohistochemistry (IHC) detection in mouse tissues, the MMTV-Cre expression pattern was examined as an indicator of PAK4 depletion. Mosaicism of Cre expression pattern in MMTV-Cre mice model has been reported ^36^. Based on estimation in both PAK4^MEp+/+^ mice and PAK4^MEp-/-^ mice, more than 90% of the mammary epithelial cells displayed Cre positive staining, and the Cre staining patterns were similar in the two groups (Fig.1d). Samples from wild-type mice without Cre expression was used here as a negative control. These results suggest that our PAK4 conditional knock-out mice have an efficient PAK4 depletion in mammary epithelial cells.

### PAK4 gene depletion in the mammary gland does not alter ductal elongation and branching

To examine the ductal growth within the mammary gland in juvenile virgin and adult virgin mice, mammary glands were isolated from mice at 4 and 10 weeks of age. To determine the potential effect of PAK4 depletion on the morphogenesis of the mammary gland, whole-mount preparations were used for measurement of the area fraction covered by mammary epithelium in the entire mammary fat pad. We found that the mammary fat pads were filled by epithelial tissue in the PAK4^MEp-/-^ mice to the same extent as in their aged matched PAK4^MEp+/+^ mice (Fig.2 a-b). We further performed a quantitative analysis of mammary tree elongation along the fat pad by measuring mammary tree length from the nipple. Quantitative analysis showed that the relative duct length in whole mounts from PAK4^MEp-/-^ mice was similar to PAK4^MEp+/+^ mice at both 4 and 10 weeks of age (Fig. 2c). To explore potential subtle differences in the ductal structure we analyzed ductal structures in hematoxylin and eosin (H&E) staining of tissue sections. This staining revealed that the ductal structures in PAK4^MEp-/-^ mammary glands were evenly distributed along the fat cells and were not distinguishable from those of PAK4^MEp+/+^ mice (Fig. 2d). Moreover, the number of mammary ducts was similar between the two groups (Fig. 2 e).

**Figure 2:**
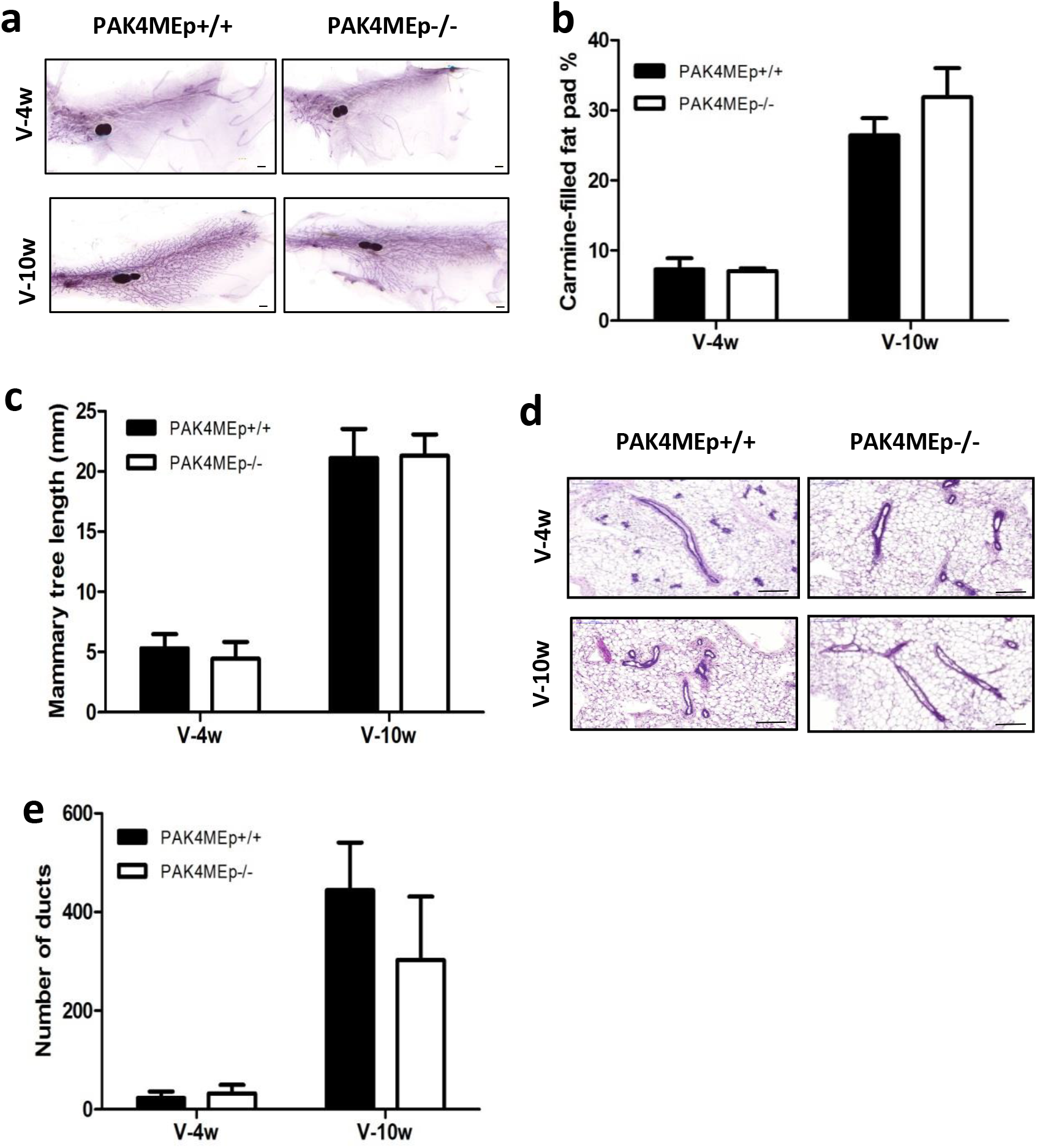
MMTV-Cre-driven PAK4 gene depletion does not impair mammary gland development. **(a)** Representative whole-mount staining of mammary glands obtained from PAK4^MEp-/-^ and PAK4^MEp+/+^ mice during the virgin week 4 and week 10 developmental stages. Scale bar is 1 mm **(b)** Whole-mount staining quantification of carmine-filled fat-pad (%) from PAK4^MEp+/+^ and PAK4^MEp-/-^ mice at virgin week 4 (P= 0.8, n=6) and week 10 (P= 0.07, n=6) stages of development. **(c)** Mammary duct length measurement in whole mounts staining from PAK4^MEp+/+^ and PAK4^MEp-/-^ mice at week 4 (P= 0.2, n=6) and week 10 (P= 0.8, n=6) of mouse development. **(d)** Representative H&E staining of mammary glands from PAK4^MEp+/+^ and PAK4^MEp-/-^ mice at week 4 (n =6) and week 10 (n =6) of mouse development. Scale bar is 200 μm. **(e)** Quantification of the number of ducts in H&E stained sections of mammary glands from PAK4^MEp-/-^ and PAK4^MEp+/+^ mice at week 4 (P= 0.9, n=6) and week 10 (P= 0.09, n=6) of mouse development. Shown values represent mean ± s.d. and p-values of unpaired t-test.

### Loss of PAK4 does not alter cell proliferation or invasion marker expression in the mammary epithelium

Considering the known role of PAK4 in cell proliferation and invasion^26,37,38^, we next sought to determine the status of cell proliferation and expression of known markers for invasion within the mammary duct in virgin week 4 mice, a time point where the mammary epithelium is highly proliferative. To quantify cell proliferation within the duct, we labeled tissues for Ki67 and consistent with our previous results, lack of Pak4 did not alter cell proliferation in PAK4^MEp-/-^ mice (Fig. 3 a-b).

**Figure 3:**
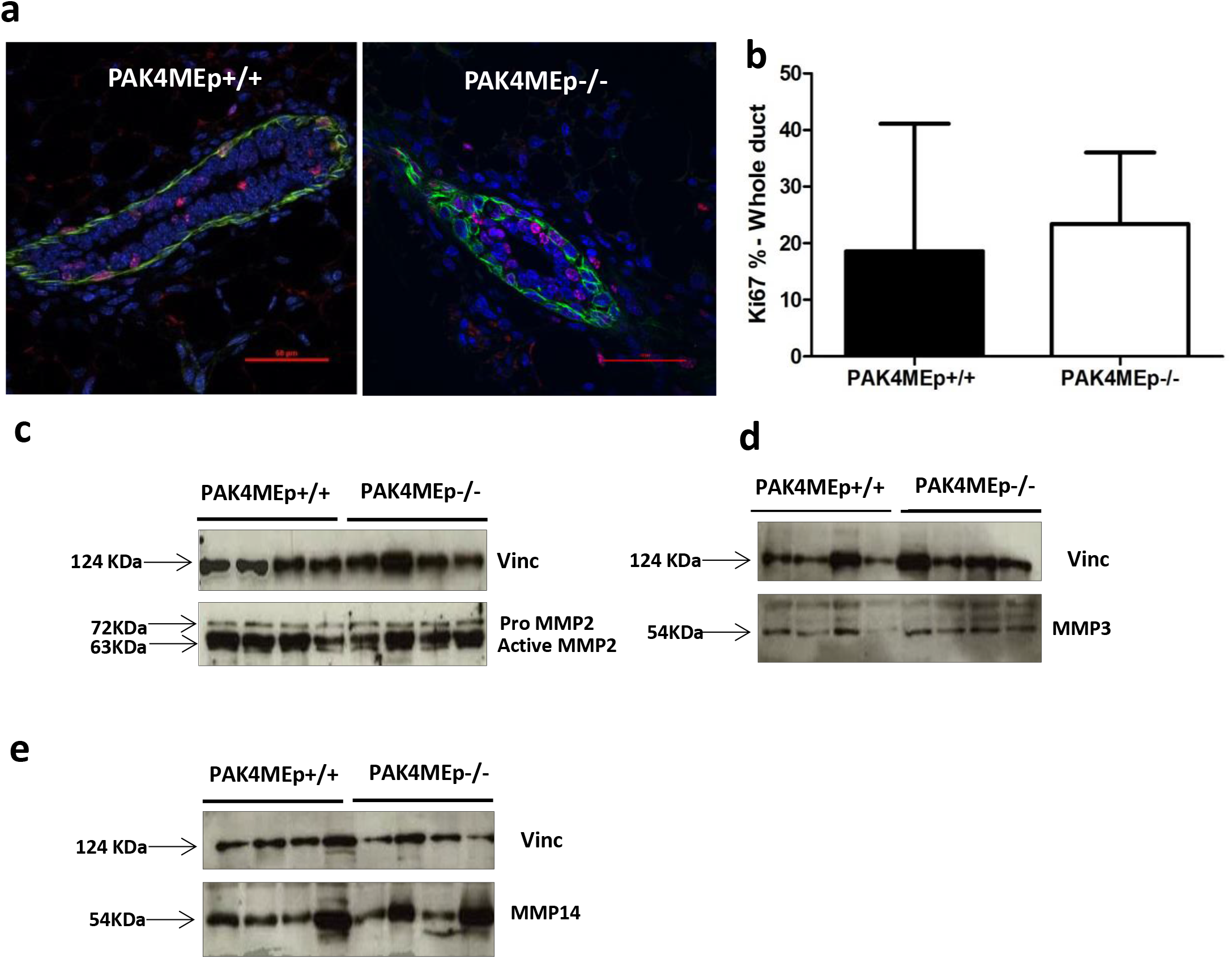
PAK4 depletion does not interfere with normal mammary epithelium proliferation. **(a)** Representative immunofluorescence labelling of Ki67 (Red) and α-SMA (Green) in inguinal mammary glands from PAK4^MEp+/+^ and PAK4^MEp-/-^ mice in virgin week 4 mice with DAPI nuclear counterstaining. **(b)** Quantification of the percentage of Ki67 positive cells in the mammary gland duct, 10 ducts from each gland were imaged and quantified. Shown values represent mean ± s.d. and p-value calculated according to unpaired t-test (P=0.6, n = 6). **(c-e)**. Immunoblot analysis of four mammary gland whole cell lysates each from PAK4^MEp+/+^ and PAK4^MEp-/-^ mice shows MMP2, MMP3 and MMP14 expressions in virgin week 4, PAK4 wild-type control and PAK4 knock-out mice. Immunoblotting for Vinculin was used as a loading control. Full gels are showed in supplementary Fig. S1b-c-d.

Next, we measured matrix metalloproteinases (MMPs) expression level, since MMPs are principal executors of matrix remodeling during mammary gland development ^39,40^. Mammary gland tissue lysates from PAK4^MEp+/+^ and PAK4^MEp-/-^ mice showed similar expression levels of MMP2, MMP3 and MMP14, suggesting no expression differences among these key invasion markers (Fig. 3 c-d-e).

### PAK4 Knock-out mothers were able to nourish pups sufficiently

We first examined and compared the overall structures of the mammary glands from PAK4 knock-out and wild-type females. In carmine red stained whole mounts from PAK4^MEp+/+^, and PAK4^MEp-/-^ mothers displayed indistinguishable ductal epithelial development at lactation day 2, and we could not detect any difference in the percentage of epithelial cells coverage of carmine filled fat pads. Moreover, the alveolar units of PAK4^MEp-/-^ females were as fully developed as in PAK4^MEp+/+^ females (Fig. 4 a-b). Both groups produced standard litter sizes (Table 1) and similar pup body weight upon weaning (Fig. 4 c). Together, this indicates that MMTV-Cre driven conditional gene depletion of PAK4 caused no defects in mammary gland development or function.

## Discussion

Using MMTV-Cre mediated PAK4 depletion in mammary gland epithelium, we have shown that targeted inactivation of PAK4 in mammary epithelial cells does not impair mammary gland development; this suggests that PAK4 is dispensable for murine mammary gland development and function.

PAK4 is ubiquitously expressed throughout embryonic development and in adult tissues ^16,41^. Transgenic mice with constitutive PAK4 gene depletion do not survive past embryonic day 11.5^16,32,33^; therefore conditional gene depletion strategies have been developed in the field to study its role in different organs development ^33^. Conditional depletion of PAK4 in heart, vasculature, and brain caused severe defect during embryonic development while PAK4 depletion in pancreas did not cause any evident defect in the pancreatic tissue development and function ^32,33,42,43^. However, the potential function of PAK4 in mammary gland development has remained unclear. To this end, we found that the mammary tree development upon PAK4 depletion was neither affected during pre-puberty nor in young adult virgin female mice; moreover PAK4 knock-out mice gave birth to healthy progeny and were able to nurse them as well as the control mice. However, it is possible that mammary gland development is dependent on other members of the PAK family, since lack of PAK1 in the mammary gland impaired lobuloalveolar development and cell differentiation ^44^, while conventional depletion of PAK5 and PAK6 resulted in normal mammary gland development ^45–48^. This supports the idea that different members of the PAK family fulfill different functions throughout organ development ^41^.

PAK4 is overexpressed in breast cancer cell lines as well as in breast cancer patients, and its overexpression is accompanied by poor patient outcome ^12,15,26^. However, our understanding of the *in vivo* function of PAK4 in breast cancer remains limited. Given that our model for conditional PAK4 gene depletion in the mouse mammary gland displays no apparent defect in organ development and function, this can serve as a useful model to study the potential *in vivo* role of PAK4 in breast cancer. Nevertheless, one should also be aware of the Cre-mosaicism that has previously been reported upon use of MMTV-Cre and similar models^35,49,50,^ which could complicate their use in an evolutionary disease like cancer, with the possibility of a selection of cells in which the gene of interest remains expressed ^50^. Using a reporter gene could be a useful approach to overcome this problem in future studies ^51^.

In summary, our data suggest that lack of PAK4 does not alter normal mammary gland development. Therefore, our mouse model of conditional depletion of PAK4 in the mammary epithelium can be useful for testing potential *in vivo* functions of PAK4 in mammary physiology and diseases such as cancer.

## Materials and Methods

### Animals

All the experimental procedures performed on animals in this study have been performed in accordance with Swedish and European Union guidelines and approved by Stockholm South and Linköping Animal Ethics Committees. To avoid the influence of social isolation, animals were housed in groups with 12:12 light: dark cycle, controlled humidity (55%± 5%), controlled temperature (21 °C ± 2 °C) and free access to food and water.

In order to generate PAK4^MEp-/-^ mice, PAK4^fl/fl^ mice (B6.129S2(FVB)-Pak4^tm2.1Amin^/J, a gift from Audrey Minden)^33^ were crossed with MMTV-Cre/Line D mice (Tg(MMTV-cre)4Mam/J, Jackson Laboratory). All mice have been maintained on a B6 background. For conditional gene depletion in the mammary gland, PAK4^fl/fl^ mice were first crossed with MMTV-Cre mice to generate MMTV-Cre; PAK4^fl/+^. Such animals were then crossed with PAK4^fl/fl^ mice, resulting in littermates with PAK^fl/fl^, PAK4^fl/+^, MMTV-Cre; PAK4^fl/+^ and MMTV-Cre; PAK4^fl/fl^ genotypes (Table 1).

Genomic DNA was prepared from biopsies using the fast tissue-to-PCR kit (#K1091, Fermentas). Mice were genotyped for heterozygous and homozygous knock-out of PAK4 according to ^33^. The primer pairs used (synthesized by ThermoFisher) were as follows: Pak4 flox: F, 5′ - CGGATATTGTCACCCACACCAG - 3′ and R, 5′ - CTAACAGGGACAGGAGCT - 3′. DNA band was visualized on 2% agar gels stained with GelRed (41003, Biotium). All mammary gland tissues used in this paper are from female mice.

### Tissue Collection

Mice were sacrificed by cervical dislocation after anesthesia with isoflurane and the mammary glands were collected. #1 and #2 thoracic mammary glands were quickly frozen and accordingly used for RNA and protein extraction. The #10 inguinal mammary gland was dissected, flattened on a piece of paper, fixed in 4% Paraformaldehyde overnight, then washed with PBS and kept in 70% ethanol for paraffin embedding and later used for immunohistochemistry.

### Whole mount staining of mammary glands

The #4 inguinal mammary gland was collected to determine the area where mammary glands were developed in fat pads. Briefly, the samples were fixed overnight with Carnoy’s fixative (100% ethanol/ chloroform /glacial acetic acid, 6:3:1). Then samples were hydrated by sequential treatment in 70%; 50%; 30%; and 10% ethanol for 15 min each. After the hydration process, samples were washed in tap water for 5 minutes and placed O/N in staining solution at RT. The staining solution was prepared by dissolving 1g carmine (C1022, Sigma) and 2.5 g aluminum potassium sulfate (A7167, Sigma) in 500 ml water followed by boiling for 20 min. The samples were then dehydrated by sequential treatment in 70%; 95%; and 100% ethanol for 15 min each, followed by storage in xylene (28975, VWR) until scanning. Scanned images were analyzed using ImageJ/Fiji (Version 1,52i) (National Institutes of Health, NIH). Whole-mount images were cropped to only include #4 inguinal mammary gland then converted to an 8-bit image and sharpened. Area of fat pad occupied by mammary epithelium (carmine staining) was measured by adjustment of threshold automatically and demonstrated as a percentage of the total mammary fat pad occupied by mammary epithelium.

Quantification of mammary tree branching was done on whole-mount images by measurement of the mammary tree length from the nipple to the last branch in millimeters using ImageJ/Fiji (Version 1,52i) (National Institutes of Health, NIH).

### H&E staining and immunostaining

Paraffin blocks were cut into 4 μm sections. For H&E staining, sections were stained with hematoxylin and eosin according to a standard protocol ^52^. For immunohistochemistry, sections were deparaffinized by incubation in 60 ° C incubator for 1 h followed by rehydration steps through washing in xylene and graded ethanol to distilled water. Samples were boiled for 20 min in antigen retrieval solution 0.01 M sodium citrate buffer (100813M, BDH) (pH 6.0) in water. Endogenous peroxidase activity was blocked via treatment with 3 % hydrogen peroxide in water (H1009, Sigma). PBS diluted normal serum from the same host species as the secondary antibody was used as a blocking buffer.

For immunohistochemistry, slides were incubated at 4 °C overnight in a humidified chamber with a α-Cre antibody (1:100, PRB-106P, Covance) diluted in blocking buffer. After three times PBS washing, slides were incubated at RT for1 h with (1:400, Biotin α-Rabbit IgG, Jackson) followed by three washes with PBS and then were incubated RT 20 min with HRP-conjugated Streptavidin (016-030-084, Jackson). Following three washes with PBS, development was performed with diaminobenzidine (DAB) (K3467, DAKO Sweden). Then slides were counterstained with Mayer HTX staining solutions (01820, Histolab), dehydrated and mounted using DPX mounting media (44581, Sigma).

For immunofluorescent experiment, sections were incubated at 4 °C ON in a humidified chamber with α-Ki67 (1:125, #12202, Santa Cruz) and α-SMA-FITC conjugated (1:300, F3777, Sigma). After washing, slides were incubated RT 1 h with (1:400, Biotin α-Rabbit IgG, Jackson) for improving signal for α-Ki67 staining. Sections were counterstained with (1:1000, Hoechst 33342, 14533, Sigma).

Images were acquired using a Nikon A1R confocal microscope with a Plan Apo VC 60×/1.4 NA oil objective and NIS-Elements software (Nikon).

### RNA extraction, reverse transcription-PCR (RT-PCR) and quantitative real-time PCR (qPCR) analysis

An RNeasy kit was utilized for RNA extraction (74104, Qiagen). RT-PCR was performed using TaqMan Reverse Transcription Reagents (N808-0234, Applied Biosystems) as described by the manufacturer. qPCR was performed as previously described^53^. qPCR was performed in a 7500 Fast Real-Time PCR System (Applied Biosystems) with FastStart Universal SYBR Green Master mix (04913914001, Roche) according to conditions specified by the manufacturer. PAK4 mRNA expression was analyzed by using primer pairs (synthesized by Invitrogen) as follows: F, 5′-GGC GCC CTC ACG GAT ATT - 3′ and R, 5′-CAC GGC GGC GAT CTG T - 3′, and the GAPDH mRNA expression was analyzed as an internal control for normalization by using following primer pairs (synthesized by Invitrogen): F, 5′-GCACAGTCAAGGCCGAGAAT-3′ and R, 5′-GCCTTCTCCATGGTGGTGAA-3′. Melting curve analysis checked the specificity of all primer pairs. All reactions were carried out in biological triplicates.

### Immunoblotting

Equal amounts of denatured protein were subjected to 10% SDS-PAGE (SDS-polyacrylamide gel electrophoresis) using molecular weight markers (26619, Fermentas) to confirm the expected size of the target proteins and transferred to PVDF membranes (IPVH00010, Millipore). Membranes were washed in TBST buffer (TBS containing 0.1% Tween-20) and non-specific binding sites were blocked by immersing the membranes in blocking buffer containing 5% non-fat milk (70166, Sigma) in TBST buffer for 1 h on a shaker at room temperature or overnight at 4°C. Membranes were first probed with the following primary antibodies: αMMP2 (1:1000, sc-10736, Santa cruz), αMMP3 (1:1000, ab52915, Abcam) and αMMP14 (1:1000, ab3644, Abcam). After the first blotting an anti-vinculin antibody (1:50000, V9131, Sigma) was used to control for protein loading. Next, peroxidase-conjugated α-rabbit (111-035-144, Jackson ImmunoResearch) secondary antibody was used. Bound antibodies were visualized with the Pierce Enhanced Chemiluminescence (ECL) Plus Western Blotting Substrate detection system (32132, ThermoFisher) according to the manufacturer’s instructions.

### Statistics

A two-tailed unpaired t-test was utilized for statistical analyses. P <0.05 was considered to represent statistical discernibility of differences.

## Author Contributions

T.Z., P.R., T.C. and S.S. designed the study; T.Z. developed the animal models, P.R., T.Z. and M.Z. collected samples; P.R. and T.Z. performed experiments and analyzed data; S.S. supervised the study and provided financial support. P.R. and S.S. drafted the paper that was then edited by all authors.

## Acknowledgments

We thank Dr. Audrey Minden for providing PAK4-floxed C57BL/6 mice. We thank Dr. Raoul Kuiper and Dr. Leander Blaas for valuable advices. This study was supported by grants to SS from the Swedish Cancer Society and the Swedish Research Council. Ting Zhuang was supported by a scholarship from the China Scholarship Council (CSC). TC was supported by the Portuguese Foundation for Science 1145 and Technology (SFRH/BD/47330/2008). Fluorescent imaging was performed at the Live Cell Imaging facility, Department of Biosciences and Nutrition, Karolinska Institutet, Sweden, supported by grants from the Knut and Alice Wallenberg Foundation, the Swedish Research Council, the Centre for Innovative Medicine and the Jonasson donation to the School of Technology and Health, Royal Institute of Technology, Sweden.

## Figure legends

**Figure 4: PAK4 depletion does not impair mammary gland lactation**

**(a)** Representative whole-mounts and H&E staining of mammary glands obtained from PAK4^MEp+/+^ and PAK4^MEp-/-^ mice in lactation day 2 stage. Scale bar is 200 μm. **(b)** Whole-mount staining quantification of carmine-filled fat-pad (%) from PAK4^MEp+/+^ and PAK4^MEp-/-^ mice at Lactation day 2(P=0.4, n=4). **(c)** Average body weights of pups from PAK4^MEp+/+^ and PAK4^MEp-/-^ mothers when weaning 21 days after birth (P=0.5, n=6). Values represent mean ± s.d. and unpaired t-test.

## Supplementary Information

**Supplementary Figure S1.**
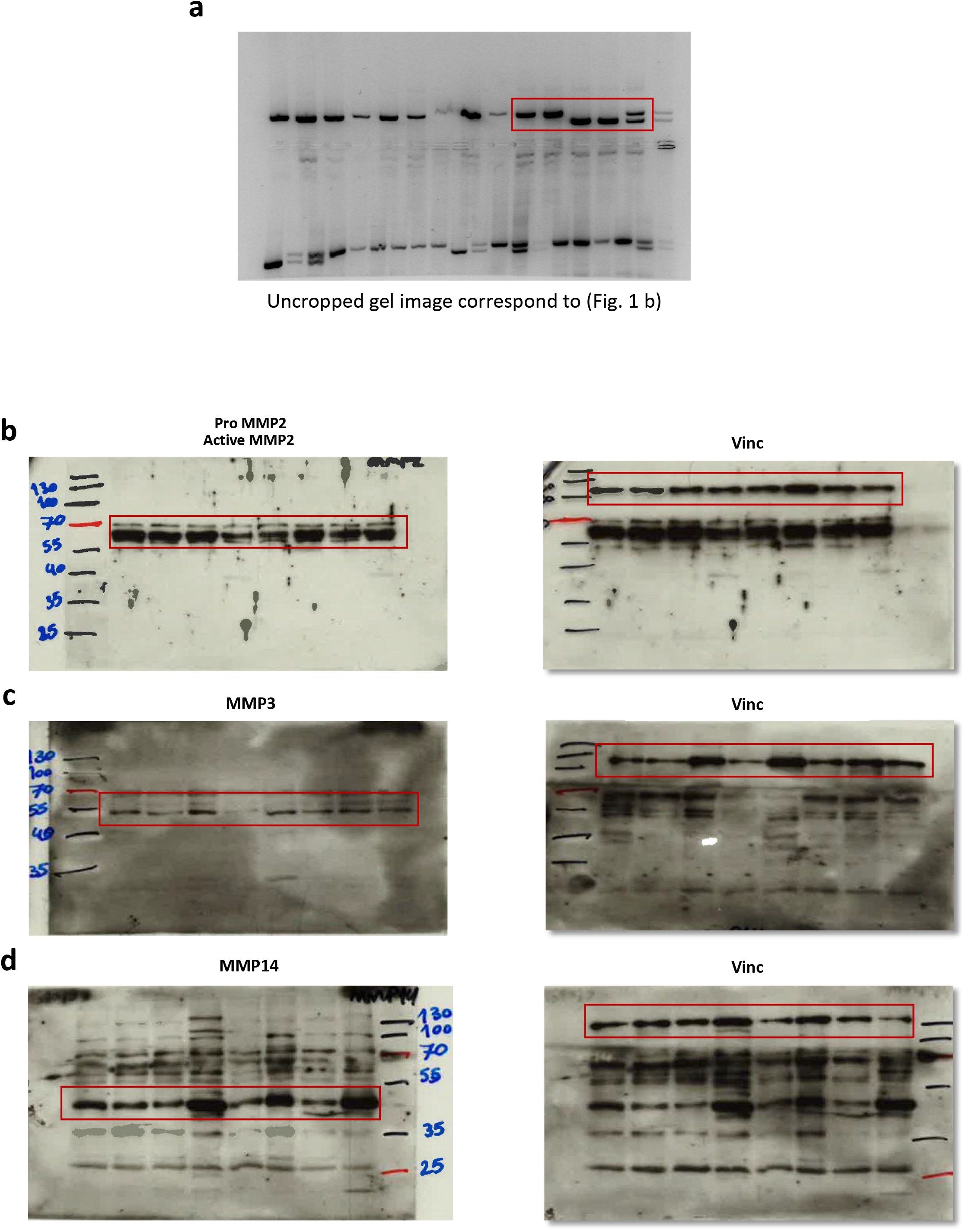
Uncropped western blot images correspond to (Fig. 3 c-d-e)

## References

1 Williams JM, D. C. Mammary ductal elongation: differentiation of myoepithelium and basal lamina during branching morphogenesis. Dev Biol 97, 274–290 (1983).

2 Lothar Hennighausen, G. W. R. Signaling Pathways in Mammary Gland Development. Developmental cell (2001).

3 Cheryll Tickle, H.-S. J. Embryonic Mammary Gland Development. eLS (2016).

4 Macias, H. & Hinck, L. Mammary gland development. Wiley Interdiscip Rev Dev Biol 1, 533–557, doi:10.1002/wdev.35 (2012).

5 Inman, J. L., Robertson, C., Mott, J. D. & Bissell, M. J. Mammary gland development: cell fate specification, stem cells and the microenvironment. Development 142, 1028–1042, doi:10.1242/dev.087643 (2015).

6 Seldin, L., Le Guelte, A. & Macara, I. G. Epithelial plasticity in the mammary gland. Curr Opin Cell Biol 49, 59–63, doi:10.1016/j.ceb.2017.11.012 (2017).

7 Kouros-Mehr, H., Slorach, E. M., Sternlicht, M. D. & Werb, Z. GATA-3 maintains the differentiation of the luminal cell fate in the mammary gland. Cell 127, 1041–1055, doi:10.1016/j.cell.2006.09.048 (2006).

8 Zaragoza, R., Garcia-Trevijano, E. R., Lluch, A., Ribas, G. & Vina, J. R. Involvement of Different networks in mammary gland involution after the pregnancy/lactation cycle: Implications in breast cancer. IUBMB Life 67, 227–238, doi:10.1002/iub.1365 (2015).

9 Katz, E. & Streuli, C. H. The extracellular matrix as an adhesion checkpoint for mammary epithelial function. Int J Biochem Cell Biol 39, 715–726, doi:10.1016/j.biocel.2006.11.004 (2007).

10 Zuo, Y., Berdeaux, R. & Frost, J. A. The RhoGEF Net1 is required for normal mammary gland development. Mol Endocrinol 28, 1948–1960, doi:10.1210/me.2014-1128 (2014).

11 Radu, M., Semenova, G., Kosoff, R. & Chernoff, J. PAK signalling during the development and progression of cancer. Nat Rev Cancer 14, 13–25 (2014).

12 Rane, C. K. & Minden, A. P21 activated kinase signaling in cancer. Semin Cancer Biol, doi:10.1016/j.semcancer.2018.01.006 (2018).

13 Itakura, A. et al. p21-Activated kinase (PAK) regulates cytoskeletal reorganization and directional migration in human neutrophils. PLoS One 8, e73063, doi:10.1371/journal.pone.0073063 (2013).

14 Kumar, R., Gururaj, A. E., & Barnes, C. J. p21-activated kinases in cancer. Nature Reviews Cancer 6, 659–471 (2006).

15 Callow, M. G. et al. Requirement for PAK4 in the anchorage-independent growth of human cancer cell lines. J Biol Chem 277, 550–558, doi:10.1074/jbc.M105732200 (2002).

16 Qu, J. et al. PAK4 kinase is essential for embryonic viability and for proper neuronal development. Mol Cell Biol 23, 7122–7133 (2003).

17 Rui-An Wang, R. K. V., Rozita Bagheri-Yarmand, Iwan Beuvink, Nancy E. Hynes, Rakesh Kumar. Essential functions of p21-activated kinase 1 in morphogenesis and differentiation of mammary glands. J Cell Biol 161, 583–592 (2003).

18 Poonam R. Molli, D.-Q. L., Murray Brion, Suresh K. Rayala, Rakesh Kumar. PAK Signaling in Oncogenesis. 28 28, 2545–2555 (2010).

19 Abo, A. et al. PAK4, a novel effector for Cdc42Hs, is implicated in the reorganization of the actin cytoskeleton and in the formation of filopodia. Embo J 17, 6527–6540, doi:10.1093/emboj/17.22.6527 (1998).

20 Jian Qu, M. S. C., Qing Shi, Kenneth C. Ha, Primal de Lanerolle, and Audrey Minden. Activated PAK4 Regulates Cell Adhesion and Anchorage-Independent Growth. Molecular and cell biology 21, 3523–3533 (2001).

21 Heckman CA, D. J., Deters D, Malwade SR, Cayer ML, Monfries C, Mamais A. Relationship of p21-activated kinase (PAK) and filopodia to persistence and oncogenic transformation. J Cell Physiol 220, 576–585 (2009).

22 Li Z, L. J., Olofsson H, Kowalewski JM, Teller S, Liu Y, Zhang H, Strömblad S. Integrin-mediated cell attachment induces a PAK4-dependent feedback loop regulating cell adhesion through modified integrin alpha v beta 5 clustering and turnover. Mol Biol Cell (2010).

23 Dart AE, B. G., Court W, Gale ME, Brown JP, Pinder SE, Eccles SA, Wells CM. PAK4 promotes kinase-independent stabilization of RhoU to modulate cell adhesion. Journal of Cell Biology 211, 863–879 (2015).

24 Zhao, M. et al. Identification of the PAK4 interactome reveals PAK4 phosphorylation of N-WASP and promotion of Arp2/3-dependent actin polymerization. Oncotarget 8, 77061–77074, doi:10.18632/oncotarget.20352 (2017).

25 Dart, A. E. & Wells, C. M. P21-activated kinase 4--not just one of the PAK. Eur J Cell Biol 92, 129–138, doi:10.1016/j.ejcb.2013.03.002 (2013).

26 Zhuang, T. et al. p21-activated kinase group II small compound inhibitor GNE-2861 perturbs estrogen receptor alpha signaling and restores tamoxifen-sensitivity in breast cancer cells. Oncotarget 6, 43853–43868, doi:10.18632/oncotarget.6081 (2015).

27 He LF, X. H., Chen M, Xian ZR, Wen XF, Chen MN, Du CW, Huang WH, Wu JD, Zhang GJ. Activated-PAK4 predicts worse prognosis in breast cancer and promotes tumorigenesis through activation of PI3K/AKT signaling. Oncotarget 14, 17573–17585 (2017).

28 Rane C, S. W., Baloglu E, Landesman Y, Crochiere M, Das-Gupta S, Minden A. A novel orally bioavailable compound KPT-9274 inhibits PAK4, and blocks triple negative breast cancer tumor growth. Sci Rep 15, 42555 (2017).

29 Hongquan Zhang, Z. L., Eva-Karin Viklund, Staffan Strömblad. p21-activated kinase 4 interacts with integrin αvβ5 and regulates αvβ5-mediated cell migration. Journal of Cell Biology 158, 1287 (2002).

30 Li Z1, Z. H., Lundin L, Thullberg M, Liu Y, Wang Y, Claesson-Welsh L, Strömblad S. p21-activated kinase 4 phosphorylation of integrin beta5 Ser-759 and Ser-762 regulates cell migration. J Biol Chem 285, 23699–23710 (2010).

31 Wagner, K. U. et al. Spatial and temporal expression of the Cre gene under the control of the MMTV-LTR in different lines of transgenic mice. Transgenic Res 10, 545–553 (2001).

32 Tian, Y., Lei, L. & Minden, A. A key role for Pak4 in proliferation and differentiation of neural progenitor cells. Dev Biol 353, 206–216, doi:10.1016/j.ydbio.2011.02.026 (2011).

33 Tian, Y., Lei, L., Cammarano, M., Nekrasova, T. & Minden, A. Essential role for the Pak4 protein kinase in extraembryonic tissue development and vessel formation. Mech Dev 126, 710–720, doi:10.1016/j.mod.2009.05.002 (2009).

34 Schwenk, F., Baron, U. & Rajewsky, K. A cre-transgenic mouse strain for the ubiquitous deletion of loxP-flanked gene segments including deletion in germ cells. Nucleic Acids Res 23, 5080–5081 (1995).

35 Wagner, K. U. et al. Cre-mediated gene deletion in the mammary gland. Nucleic Acids Res 25, 4323–4330 (1997).

36 Robinson, G. W. & Hennighausen, L. MMTV-Cre transgenes can adversely affect lactation: considerations for conditional gene deletion in mammary tissue. Anal Biochem 412, 92–95, doi:10.1016/j.ab.2011.01.020 (2011).

37 Siu, M. K. et al. p21-activated kinase 4 regulates ovarian cancer cell proliferation, migration, and invasion and contributes to poor prognosis in patients. Proc Natl Acad Sci USA 107, 18622–18627, doi:10.1073/pnas.0907481107 (2010).

38 Close & D. Kesanakurti, C. C., D. Rajasekhar Maddirela, M. Gujrati, J.S. Rao. Functional cooperativity by direct interaction between PAK4 and MMP-2 in the regulation of anoikis resistance, migration and invasion in glioma. Cell death Disease 3, 445 (2012).

39 Talhouk, R. S., Chin, J. R., Unemori, E. N., Werb, Z. & Bissell, M. J. Proteinases of the mammary gland: developmental regulation in vivo and vectorial secretion in culture. Development 112, 439–449 (1991).

40 Correia AL, M. H., Chen EI, Schmitt FC, Bissell MJ. The hemopexin domain of MMP3 is responsible for mammary epithelial invasion and morphogenesis through extracellular interaction with HSP90β. Genes Dev 27, 805–817 (2013).

41 Kelly, M. L. & Chernoff, J. Mouse models of PAK function. Cell Logist 2, 84–88, doi:10.4161/cl.21381 (2012).

42 Nekrasova T, M. A. Role for p21-activated kinase PAK4 in development of the mammalian heart. Transgenic Res 21, 797–811 (2012).

43 Zhao, M. et al. Pdx1-Cre-driven conditional gene depletion suggests PAK4 as dispensable for mouse pancreas development. Sci Rep 7, 7031, doi:10.1038/s41598-017-07322-5 (2017).

44 Rui-An Wang, R. K. V., Rozita Bagheri-Yarmand, Iwan Beuvink, Nancy E. Hynes, and Rakesh Kumar. Essential functions of p21-activated kinase 1 in morphogenesis and differentiation of mammary glands. J Cell Biol 161, 583 (2003).

45 Furnari, M. A., Jobes, M. L., Nekrasova, T., Minden, A. & Wagner, G. C. Differential sensitivity of Pak5, Pak6, and Pak5/Pak6 double-knockout mice to the stimulant effects of amphetamine and exercise-induced alterations in body weight. Nutritional neuroscience 17, 109–115 (2014).

46 Furnari, M. A., Jobes, M. L., Nekrasova, T., Minden, A. & Wagner, G. C. Functional deficits in PAK5, PAK6 and PAK5/PAK6 knockout mice. PLoS One 8, e61321 (2013).

47 Nekrasova, T., Jobes, M. L., Ting, J. H., Wagner, G. C. & Minden, A. Targeted disruption of the Pak5 and Pak6 genes in mice leads to deficits in learning and locomotion. Dev Biol 332, 95–108 (2008).

48 Li, X. M., A. Targeted disruption of the gene for the PAK5 kinase in mice. Mol Cell Bio 23, 7134–7142 (2003).

49 Wagner KU, M. K., Ward T, Davis B, Wiseman R, Hennighausen L Spatial and temporal expression of the Cre gene under the control of the MMTV-LTR in different lines of transgenic mice. Transgenic Res 10, 545–553 (2001).

50 Gertraud W Robinson, L. H. MMTV-Cre transgenes can adversely affect lactation: considerations for conditional gene deletion in mammary tissue. Anal Biochem 412, 92–95 (2011).

51 Sakamoto K, S. J., Wagner K-U. Generation of a Novel MMTV-tTA Transgenic Mouse Strain for the Targeted Expression of Genes in the Embryonic and Postnatal Mammary Gland. PLoS One 7, e43778 (2012).

52 Cardiff, R. D., Miller, C. H. & Munn, R. J. Manual hematoxylin and eosin staining of mouse tissue sections. Cold Spring Harb Protoc 2014, 655–658, doi:10.1101/pdb.prot073411 (2014).

53 Qiao, Y. et al. AP-1-mediated chromatin looping regulates ZEB2 transcription: new insights into TNFalpha-induced epithelial-mesenchymal transition in triple-negative breast cancer. Oncotarget 6, 7804–7814, doi:10.18632/oncotarget.3158 (2015).

